# Nuclear Cathepsin L Remodels the Replication Machinery to Create a Therapeutic Vulnerability in Ovarian Cancer

**DOI:** 10.64898/2026.09.02.748933

**Authors:** Prabhu Thirusangu, Ling Jin, Shruti Rao, Karthik Balakrishnan, Xiaonan Hou, Jamison L VanBlaricom, Sun-Hee Lee, Ann L Oberg, John Weroha, Scott H Kaufmann, Jamie N Bakkum-Gamez, Viji Shridhar

## Abstract

Therapeutic resistance in ovarian cancer is frequently driven by persistent replication stress, yet the molecular mechanisms that convert replication stress into a therapeutically exploitable vulnerability remain incompletely understood. Here, we identify drug-induced nuclear cathepsin L (nCTSL) as a previously unrecognized regulator of replication stress and DNA repair. Clofarabine (CLF) combined with the ATR inhibitor AZD6738, or the CHK1 inhibitor prexasertib promoted nuclear accumulation of CTSL, where it remodeled the replication machinery through degradation of CCNE1, MCM3, MCM6, and geminin, accompanied by loss of RAD51 and 53BP1 and increased γH2AX and phospho-RPA2. DNA fiber analysis demonstrated marked inhibition of replication fork progression following CLF-based combinations, whereas CTSL depletion accelerated fork progression and abolished therapy-induced replication stress. Reconstitution with the nuclear M1F CTSL isoform restored replication restraint, confirming a direct role for nuclear CTSL in regulating replication dynamics. GFP-based DNA repair reporter assays further revealed that CLF-based combinations suppress DNA repair competence in a CTSL-dependent manner, indicating that nuclear CTSL couples replication stress amplification with functional inhibition of repair pathways. Functionally, CLF-based combinations selectively targeted transformed fallopian tube secretory epithelial cells while sparing non-transformed counterparts, demonstrated broad activity in patient-derived ovarian cancer ascites spheroids, and significantly inhibited tumor growth and prolonged survival *in vivo*. Collectively, our findings identify nuclear CTSL as a mechanistic driver of replication stress that remodels the replication machinery, impairs DNA repair, and creates a therapeutically exploitable vulnerability in ovarian cancer. We propose that nuclear CTSL promotes a transition from replication competence to replication catastrophe, thereby establishing a conceptual framework for biomarker-guided therapeutic strategies targeting CTSL-dependent replication stress.

## INTRODUCTION

High-grade serous ovarian cancer (HGSOC) is characterized by extensive genomic instability and persistent replication stress arising from TP53 loss, oncogene activation, and uncontrolled replication origin firing [1, 2]. Although this chronic replication stress promotes tumor evolution and therapeutic resistance, it also creates a potential vulnerability because cancer cells become highly dependent on replication stress response (RSR) pathways to maintain DNA replication and genome integrity. Consequently, inhibitors of ATR, CHK1, and PARP have emerged as important therapeutic strategies for exploiting replication stress in ovarian cancer [3–5]. Despite encouraging preclinical activity, however, clinical responses remain heterogeneous and are frequently limited by intrinsic or acquired resistance, highlighting an incomplete understanding of the molecular events that determine whether replication stress remains compatible with survival or progresses to irreversible replication collapse.

Current therapeutic strategies primarily target signaling pathways that stabilize stalled replication forks or coordinate DNA repair. However, considerably less is known about how the replication machinery itself is remodeled during therapeutic stress. Replication competence is maintained through the coordinated activity of origin licensing factors, replicative helicases, cell-cycle regulators, and homologous recombination machinery, which collectively ensure accurate DNA synthesis, replication fork progression, and genome stability [6–8]. Whether selective disruption of this machinery can convert replication-competent cancer cells into a state of replication collapse remains largely unknown.

Cathepsin L (CTSL) is a lysosomal cysteine protease increasingly recognized to possess important nuclear functions [9–11]. Alternative translation initiation generates a signal peptide-deficient CTSL isoform capable of nuclear localization, where CTSL has been implicated in chromatin remodeling [12, 13], transcriptional regulation, and DNA repair [10, 13, 14]. We recently demonstrated that clofarabine promotes KPNB1-dependent nuclear accumulation of CTSL, resulting in degradation of RAD51 and 53BP1 and sensitization of ovarian cancer cells to PARP inhibition [15]. These findings suggested that nuclear CTSL functions as an active regulator of genome maintenance rather than simply a lysosomal protease.

Whether nuclear CTSL directly regulates replication machinery and the functional consequences of this activity have remained unknown. We hypothesized that therapeutic induction of nuclear CTSL proteolytically remodels key components of the replication machinery, compromising replication competence, impairing DNA repair, and creating a therapeutically exploitable vulnerability in ovarian cancer.

Here, we show that combinations of clofarabine with PARP, ATR, or CHK1 inhibition promote nuclear accumulation of CTSL, where it degrades multiple regulators of replication licensing, fork progression, and DNA repair, including CCNE1, MCM3, MCM6, geminin, RAD51, and 53BP1. These changes functionally impair replication fork progression, promote replication collapse, inhibits repair and produce robust therapeutic responses in ovarian cancer cell lines, patient-derived ascites spheroids, and in vivo models. Together, our findings identify nuclear CTSL as a previously unrecognized regulator of replication competence that remodels the replication machinery to create a therapeutically exploitable vulnerability in ovarian cancer.

## RESULTS

### Drug-induced nuclear CTSL selectively creates a therapeutic vulnerability in transformed fallopian tube secretory epithelial cells

To determine whether CLF-based therapeutic combinations selectively target transformed epithelial cells while sparing their non-transformed counterparts, we compared the effects of CLF combined with the CHK1 inhibitor prexasertib or the ATR inhibitor ceralasertib (AZD6738) in a genetically defined series of human fallopian tube secretory epithelial cells (FTSECs). These included immortalized FT33 cells generated by expression of hTERT and SV40 large T antigen, together with matched transformed derivatives generated by expression of mutant CDK4 (CDK4R24C) and shRNA-mediated p53 suppression or by c-Myc overexpression [16, 17].

Clonogenic survival assays demonstrated that FT33 cells remained highly resistant to prexasertib at concentrations up to 2 nM, either alone or in combination with 1 nM CLF (Fig. 1A). In contrast, both transformed FT33-shP53/CDK4R24C cells and tumorigenic FT33-Myc cells exhibited marked sensitivity to prexasertib, which was further enhanced by CLF treatment (Fig. 1B,C). Addition of CLF shifted the prexasertib dose-response curve approximately two-fold, reducing the IC50 from approximately 1.0 nM to 0.5 nM in both transformed models. Similar selectivity was observed following combined inhibition of ATR, where CLF plus AZD6738 effectively suppressed colony formation in transformed FTSECs but had minimal effects on immortalized FT33 cells (Figure S1). Collectively, these findings demonstrate that CLF-based combinations preferentially target transformed ovarian epithelial cells while sparing non-transformed FTSECs, indicating the existence of a therapeutically exploitable window associated with malignant transformation.

**Figure 1.**
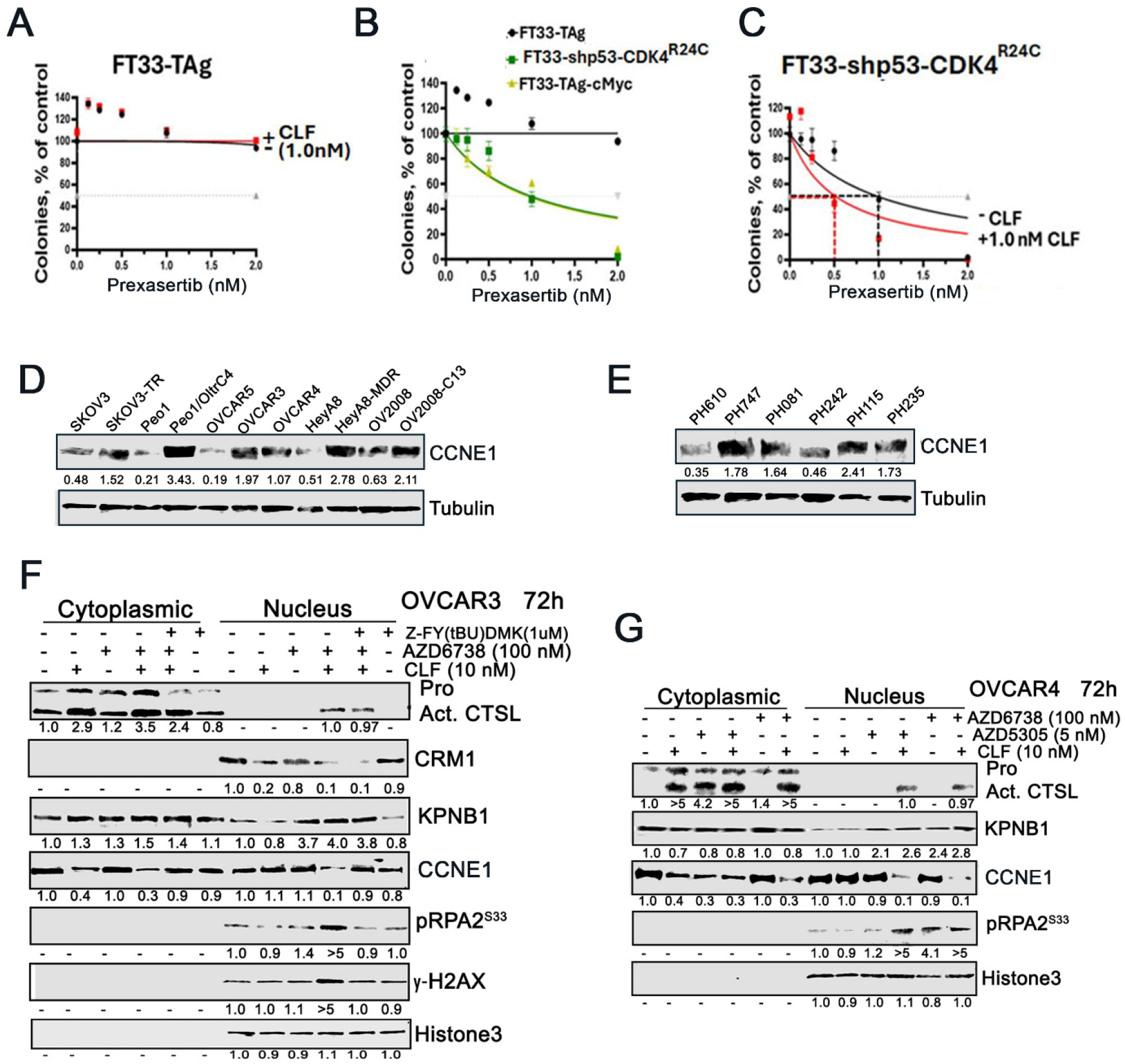
Inhibition of PARP and ATR/Chk1 pathway facilitates the CLF to induce nuclear CTSL in CCNE1 amplified OC cells. **A)** Colony formation assays showing that immortalized FT33 cells (hTERT + SV40 T antigen) are resistant to prexasertib (≤ 2 nM), both with and without CLF (1.0 nM). **B, C)** Transformed FT33-shP53/CDK4R24C and **C)** FT33 c-Myc/TAg cells display marked sensitivity to prexasertib monotherapy (black lines). Addition of CLF (1.0 nM) further shifts the prexasertib dose–response curve (red lines), enhancing sensitivity and reducing the prexasertib IC₅₀ approximately two-fold (from ∼1.0 nM to ∼0.5 nM). These data indicate selective synergy between CLF and prexasertib in transformed, but not non-transformed, fallopian tube epithelial cells. **D, E)** Immunoblot analysis demonstrates elevated CCNE1 (Cyclin E1) expression in drug-resistant isogenic cell lines relative to parental counterparts and *CCNE1*-amplified (PH115, PH235) or PARPi-resistant ovarian cancer PDX models., respectively. **F)** Western blot analysis of CTSL, KPNB1, CRM1, Cyclin E1, pRPA2, and *γγ*H2AX in cytoplasmic and nuclear fractions of *CCNE1*-amplified OVCAR3 cells treated with indicated drugs ± CTSL inhibitor for 72 hours. Fold changes relative to untreated controls were normalized using ImageJ software. **G)** Similar subcellular fractionation and western blot analysis in OVCAR4 cells following 72 hours treatment with indicated drugs.

### Nuclear CTSL proteolytically remodels the replication machinery

Since transformed tumors frequently depend upon elevated replication stress for continued proliferation, we investigated whether nuclear CTSL regulates proteins governing replication competence. *CyclinE1* (CCNE1) amplification (AMP) or overexpression (OE), which is found in 20-40% [18] of OC cases, is strongly linked to HR proficiency, chemoresistance, and poor outcomes and is a marker of replication stress[19]. Western blot analysis demonstrated that CCNE1 expression was markedly elevated in multiple drug-resistant ovarian cancer models, including resistant isogenic derivatives and patient-derived xenograft (PDX) cultures exhibiting CCNE1 amplification and homologous recombination proficiency (Fig. 1D, E). These observations are consistent with the established role of CCNE1 amplification in promoting replication origin firing and therapeutic resistance and suggested that CCNE1-high tumors [20]may provide an appropriate context for evaluating CTSL-dependent regulation of the replication machinery. Subcellular fractionation demonstrated that treatment with CLF plus AZD6738 induced robust nuclear accumulation of activated CTSL in CCNE1-high OVCAR3 and OVCAR4 cells (Fig. 1F, G). Nuclear CTSL accumulation closely coincided with phosphorylation of RPA32^Ser33^ and accumulation of γH2AX, indicating activation of replication-associated DNA damage signaling. Importantly, these molecular changes occurred concomitantly with marked attenuation of nuclear CCNE1, whereas treatment with the CTSL inhibitor Z-FY-(tBu)DMK preserved CCNE1 expression and substantially reduced both pRPA2^Ser33^ and γH2AX despite identical drug exposure. These findings demonstrate that CTSL proteolytic activity is required for efficient CCNE1 loss and the accompanying replication stress response.

Mechanistically, CLF treatment reduced expression of the nuclear export receptor CRM1 while promoting KPNB1-dependent nuclear retention of CTSL, thereby favoring sustained nuclear accumulation of the protease[15]. Although ATR inhibition alone induced modest pRPA2^Ser33^ activation, this occurred in the absence of detectable nuclear CTSL accumulation and without significant CCNE1 downregulation. In contrast, combined CLF and ATR inhibition produced substantially greater activation of replication stress signaling that directly correlated with persistent nuclear CTSL accumulation and loss of CCNE1. Similar CTSL-dependent regulation was observed following CLF combined with the PARP inhibitor saruparib (AZD5305), indicating that induction of nuclear CTSL represents a common response to mechanistically distinct CLF-based therapeutic combinations (Fig. 1G).

To further determine whether CCNE1 attenuation reflected a broader effect on the replication machinery, we examined additional proteins required for replication licensing and replisome function. In CCNE1-high PH081 patient-derived ovarian cancer cells, treatment with clinically relevant concentrations of CLF and prexasertib induced robust nuclear accumulation of CTSL together with coordinated reduction of CCNE1, geminin, MCM3, and MCM6 (Fig. 2A). These molecular changes occurred simultaneously with increased pRPA2^Ser33^ and γH2AX, indicating progressive disruption of replication-associated homeostasis. Importantly, pharmacologic inhibition of CTSL preserved expression of CCNE1, geminin, and MCM3 while markedly reducing activation of replication stress signaling, despite continued inhibition of CHK1 phosphorylation by prexasertib. Comparable CTSL-dependent remodeling of replication-associated proteins was observed in additional ovarian cancer models, including PEO1 and CCNE1-overexpressing mouse KPCA cells [21] (Fig. 2B, C), demonstrating that this response is reproducible across genetically diverse ovarian cancer models. Rather than selectively targeting individual proteins, nuclear CTSL coordinately disrupted multiple functional components required for replication licensing, replisome integrity, and cell-cycle progression. Collectively, these findings identify nuclear CTSL as a proteolytic regulator of the replication machinery that remodels the molecular framework required to maintain replication competence.

**Figure 2.**
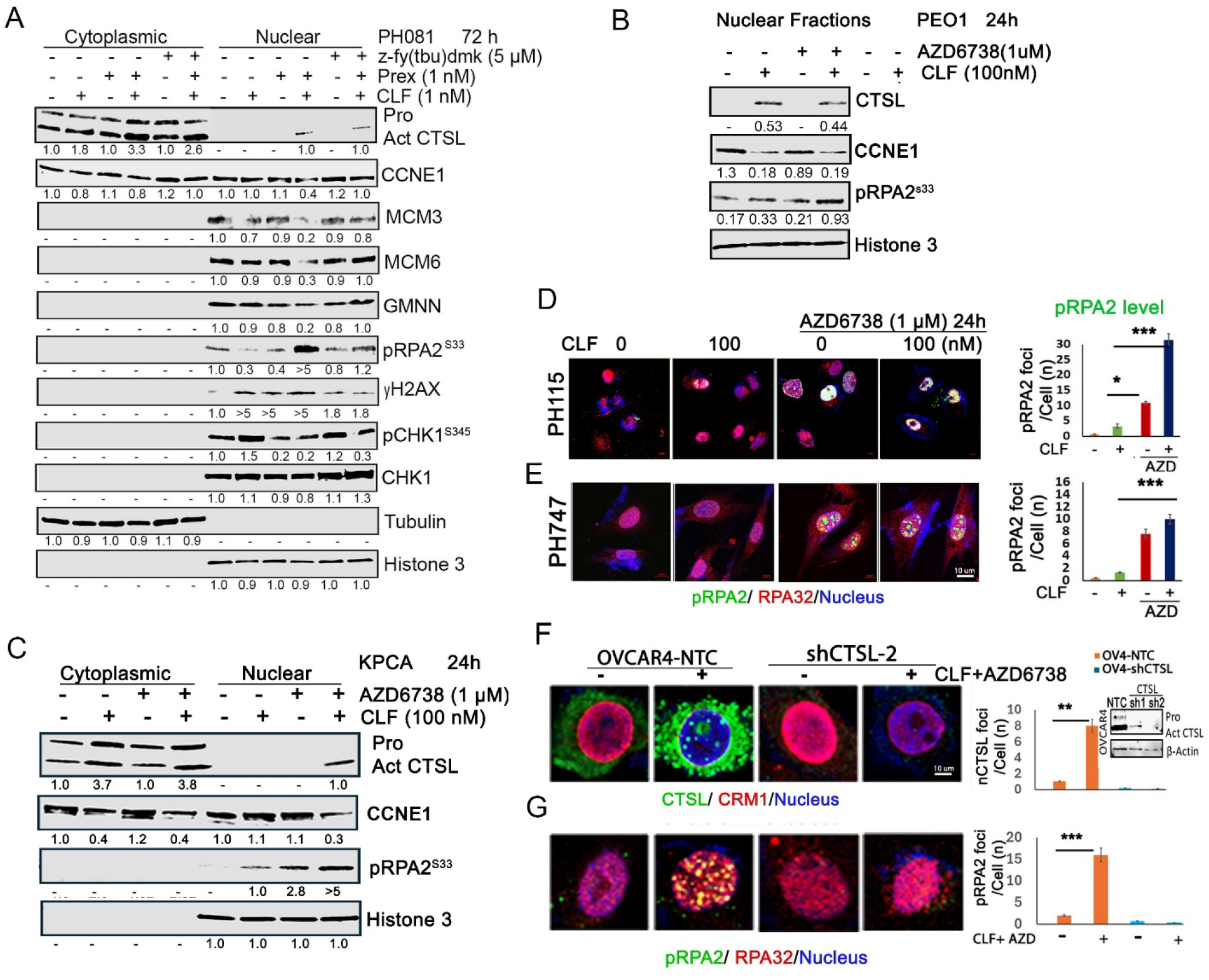
Nuclear CTSL drives loss of replication licensing factors and enforces replication stress. **A)** Western blot analysis of CTSL, CCNE1, MCM3, MCM6, GMNN, pRPA2, γH2AX, pChk1, and total Chk1 (tChk1) levels in cytoplasmic and nuclear fractions from PH081 cultures treated with the indicated drugs ± CTSL inhibitor for 72 hours. Fold changes relative to untreated controls were normalized using ImageJ software. **B)** Immunoblot analysis of CTSL, CCNE1, and pRPA2 in nuclear fractions from PEO1 cells treated with CLF and AZD6738 at indicated concentrations for 24 hours. **C)** Similar subcellular fractionation and western blot analysis of CCNE1-overexpressing KPCA cells, with fold changes normalized to untreated controls via ImageJ. **D, E)** Immunofluorescence staining of pRPA2 (S33) foci in *ex vivo* PDX cultures PH115 and PH747 respectively, following treatment with indicated concentrations of CLF and AZD6738. Representative bar graphs display the number of pRPA2 foci per cell. **F)** Immunofluorescence analysis of nuclear CTSL (nCTSL) foci and CRM1 levels in OVCAR4 cells treated with CLF and AZD6738 at indicated concentrations. The bar graph shows nCTSL foci counts per cell, and the immunoblot insert confirms stable CTSL knockdown in OVCAR4 cells. **G)** Immunofluorescence analysis of pRPA2 foci under similar treatment conditions in OVCAR4 (NTC, shCTSL-2) cells, with the representing bar graph displaying pRPA2 foci counts per cell. α-Tubulin and Histone H3 served as cytoplasmic and nuclear loading controls, respectively. Statistics were analyzed by students t-test and significances were expressed as *\*p < 0.05, **p < 0.01 and ***p < 0.001*.

### Nuclear CTSL-mediated remodeling of the replication machinery compromises DNA repair and replication stress signaling

Having established that nuclear CTSL coordinately remodels proteins governing replication licensing and replisome integrity, we next determined whether these molecular alterations functionally impair DNA repair pathways required to maintain replication fork stability. Because RAD51 and 53BP1 are central regulators of homologous recombination and DNA damage repair, and were previously identified as CTSL substrates, we investigated whether CLF-based therapeutic combinations further compromise repair capacity by simultaneously disrupting replication machinery and DNA repair proteins. Immunofluorescence analysis of CCNE1-amplified patient-derived ovarian cancer cultures (PH115 and PH747) demonstrated that treatment with CLF plus ceralasertib produced a marked increase in pRPA2^Ser33^ nuclear foci compared with either agent alone (Fig. 2D, E). Quantitative analysis confirmed a significant increase in replication stress-associated pRPA2^Ser33^ foci following combination treatment, consistent with activation of replication-associated DNA damage signaling in clinically relevant patient-derived models.

To determine whether this response required CTSL, OVCAR4 cells expressing non-targeting shRNA (NTC) or CTSL-specific shRNA were examined following CLF plus ATR inhibition. Combination treatment induced robust nuclear accumulation of CTSL in control cells, which coincided with abundant pRPA2^Ser33^ foci. In contrast, stable CTSL knockdown almost completely abolished induction of pRPA2^Ser33^ despite identical drug exposure, demonstrating that activation of replication stress signaling requires nuclear CTSL (Fig. 2F, E). Similar findings were obtained following CLF combined with prexasertib, where CTSL depletion prevented induction of both pRPA2^Ser33^ and γH2AX foci (Fig. S2), further demonstrating that nuclear CTSL functions upstream of replication-associated DNA damage signaling irrespective of whether replication stress is potentiated through ATR or CHK1 inhibition. Because checkpoint activation represents a critical adaptive response that preserves replication fork integrity, we next examined the contribution of ATR signaling to CLF sensitivity. Stable depletion of ATR, but not ATM, markedly sensitized PEO1/OLTRC4 ovarian cancer cells to CLF monotherapy, shifting CLF sensitivity from the micromolar to nanomolar range (Fig. 3A, B). Consistent with these findings, combining low-dose CLF with AZD6738 produced pronounced synergistic inhibition of clonogenic survival in both PEO1/OLTRC4 and OVCAR3 cells (Fig. 3C, D).

**Figure 3.**
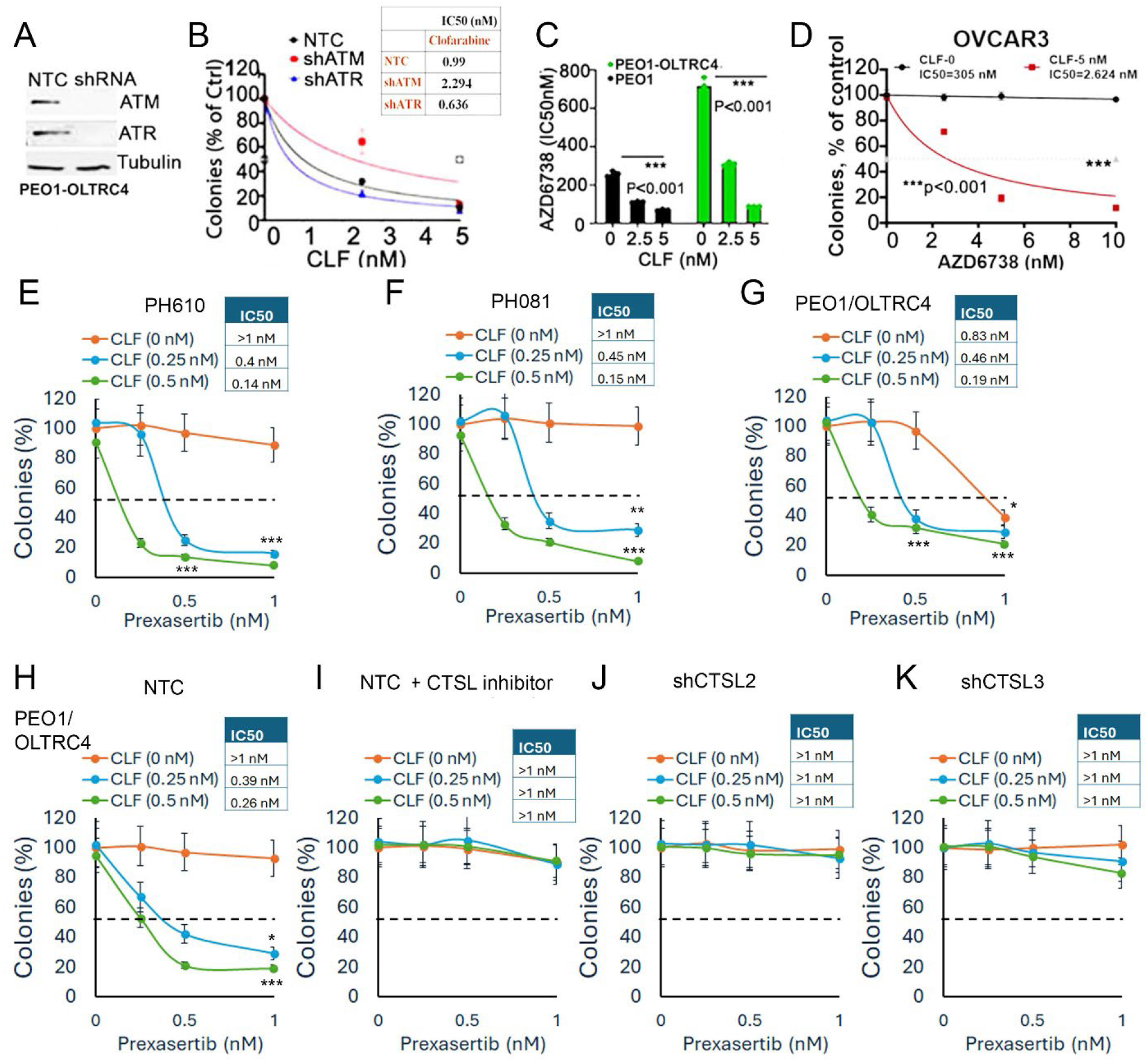
Genetic and pharmacological inhibition of ATR or Chk1 enhances sensitivity of OC cells to CLF treatment. A) Western blot analysis confirming stable knockdown of ATR and ATM in PEO1/OLTRC4 cells. B) Colony formation assays in PEO1/OLTRC4 cells demonstrating that ATR knockdown, but not ATM knockdown, sensitizes cells to CLF monotherapy and improves IC₅₀ values. C) Combination treatment with CLF (2.5–5.0 nM) and AZD6738 further enhances sensitivity by ∼3- to 6-fold in PEO1/OLTRC4 cells. D) Clonogenic assays in OVCAR3 cells showing the synergistic effect of CLF and AZD6738 combinations at the indicated concentrations. **E–G)** Colony formation assays demonstrating strong synergy between CLF and prexasertib in e*x vivo* CCNE1-amplified PDX models PH610, PH081 and cell line PEO1/OLTRC4 respectively, showing enhanced prexasertib potency and improved IC₅₀ in the presence of CLF. **H, I)** Synergy assays in NTC-control PEO1/OLTRC4 cells showing that CLF + prexasertib combination induced cytotoxicity is rescued in the presence of a CTSL-specific inhibitor Z-FY(tBU)DMK (5 μM). **J, K)** Similar colony formation assays in CTSL-knockdown cells (sh2, sh3) demonstrating resistance to the CLF + prexasertib combination. Statistical significance was determined using Student’s t-test (p < 0.05, *p < 0.01, p < 0.001).

Since prexasertib directly targets CHK1, we next evaluated whether CHK1 inhibition similarly cooperates with CLF to exploit CTSL-dependent remodeling of replication competence. Across multiple ovarian cancer models, including PH610, PH081, and PEO1/OLTRC4 cells, CLF and prexasertib produced robust synergy, reducing the prexasertib IC_50_ approximately six- to seven-fold while using physiologically relevant concentrations of CLF (Fig. 3E-G). To directly establish the requirement for CTSL in mediating this response, PEO1/OLTRC4 cells expressing either non-targeting shRNA or independent CTSL shRNAs were examined. Whereas control cells exhibited marked sensitivity to CLF plus prexasertib, pharmacologic inhibition of CTSL or stable CTSL knockdown substantially attenuated therapeutic synergy and restored clonogenic survival (Fig. 3H - K). Together, these findings establish that nuclear CTSL is required for the enhanced therapeutic efficacy observed following combined CLF and checkpoint inhibition. Collectively, these experiments demonstrate that nuclear CTSL-mediated remodeling of the replication machinery extends beyond disruption of replication licensing proteins to functionally compromise homologous recombination, amplify replication-associated DNA damage signaling, and diminish checkpoint-dependent recovery pathways. These coordinated molecular changes predict that cells undergoing CTSL-dependent remodeling should progressively lose the capacity to sustain productive DNA replication.

### Nuclear CTSL-mediated remodeling functionally impairs replication fork progression

To determine whether remodeling of the replication machinery produces functional defects in DNA replication, we directly examined replication fork dynamics using single-molecule DNA fiber analysis. Cells were sequentially labeled with CldU and IdU following treatment with physiologically relevant concentrations of CLF and prexasertib, either alone or in combination, allowing direct measurement of replication fork progression. Neither CLF nor prexasertib alone produced substantial impairment of replication fork progression under these conditions despite their established effects on nucleotide metabolism and checkpoint signaling (Fig. 4A). In contrast, combined treatment with CLF and prexasertib caused a profound reduction in IdU tract length, indicating marked inhibition of ongoing DNA synthesis and replication fork progression. These findings demonstrate that simultaneous induction of nuclear CTSL together with checkpoint inhibition produces functional impairment of DNA replication that exceeds the effects of either agent alone.

**Figure 4.**
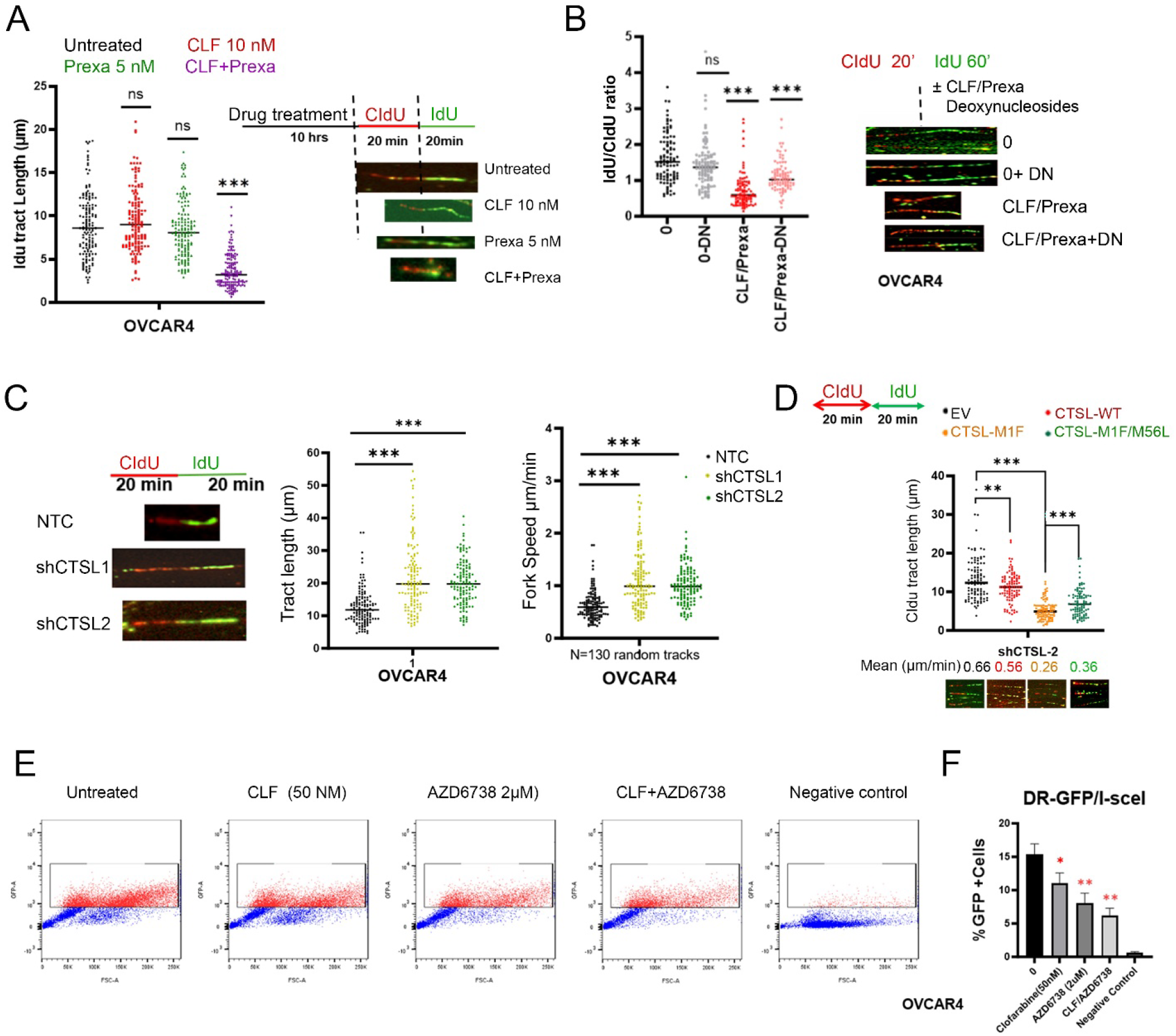
The effect of clofarabine and prexasertib and nCTSL on replication fork progress at single molecule DNA fiber. **A)** DNA fiber analysis of OVCAR4 cells treated with CLF and prexasertib at the indicated concentrations for 10 hours, followed by sequential labeling with CldU (20 min) and IdU (20 min). Representative images of DNA fibers and quantification of IdU tract lengths are shown. **B)** DNA fiber analysis of OVCAR4 cells labeled with CldU (20 min) and then relabeled with IdU (60 min) in the presence of the CLF/prexasertib combination with or without exogenous deoxynucleosides (10 µM). Representative images of DNA fibers and quantification of the IdU/CldU tract length ratios are shown. **C, D)** DNA fiber analysis performed in C) CTSL-knockdown OVCAR4 cells (sh1, sh2) and D) OVCAR4 shCTSL-2 cells ectopically overexpressing CTSL-WT, CTSL-M1F, or CTSL-M1F/M56. Representative images of DNA fibers alongside quantifications of IdU tract length and replication fork speed (*n* = 153) are shown. Bars represent medians. **E)** Representative flow cytometry plots showing the percentage of GFP-positive OVCAR4 cells as a measure of double-strand break (DSB) repair following a 24-hour treatment with the indicated concentrations of CLF and AZD6738, alone or in combination. **F)** Quantification of GFP-positive cells presented as mean ± SD. Statistical significance was determined using a Kruskal-Wallis test with Dunn’s post-hoc analysis (*p* < 0.05, \**p* < 0.01, *p* < 0.001; ns = non-significant) compared to untreated controls or between indicated groups.

Because IdU incorporation following combination treatment was markedly reduced, the duration of the second labeling pulse was extended from 20 minutes to 60 minutes to determine whether impaired fork progression reflected delayed DNA synthesis rather than complete loss of replication competence. Although prolonged labeling modestly increased IdU incorporation, replication tracts remained substantially shorter than untreated controls, confirming persistent inhibition of replication fork progression (Fig. 4B). Supplementation with exogenous deoxynucleosides partially restored IdU tract length following CLF or prexasertib monotherapy, indicating that depletion of nucleotide pools contributes to impaired replication under these conditions. However, deoxynucleoside supplementation failed to fully restore replication following combined treatment, demonstrating that impaired fork progression cannot be explained solely by nucleotide depletion and instead reflects additional disruption of the replication machinery.

To determine whether endogenous CTSL contributes to basal regulation of DNA replication, we performed DNA fiber analysis in OVCAR4 cells following stable CTSL depletion. Sequential CldU/IdU labeling revealed a significant increase in replication fork speed in both independent CTSL knockdown lines compared with non-targeting control cells (Fig. 4C). Whereas control cells exhibited relatively restrained fork progression, depletion of CTSL markedly accelerated replication dynamics, indicating that endogenous CTSL functions as a negative regulator of replication fork progression under basal conditions. These findings demonstrate that CTSL is required for maintenance of replication restraint and suggest that loss of CTSL promotes a replication-competent state characterized by enhanced fork progression. Further quantification of 130 random replication tracks demonstrates that CTSL depletion increases median fork speed from approximately 0.5–0.6 μm/min in NTC cells to approximately 1.0 μm/min in CTSL-deficient cells (Fig. 4C), representing nearly a two-fold increase in replication velocity. Recently we reported that ectopic expression wild type CTSL and the CTSL mutated at methionine 1 (M1F) translocated into nucleus and however, the CTSL mutated at methionine 1 and 56 (M1F/M56L) fails to translocate into the nucleus [15]. In the current study, to directly determine whether nuclear CTSL itself is sufficient to impair DNA replication, DNA fiber analysis was performed in CTSL-depleted OVCAR4 cells reconstituted with wild-type CTSL, the signal peptide-deficient nuclear M1F isoform, or the trafficking-defective M56L mutant. Whereas vector control and wild-type CTSL cells exhibited relatively preserved replication dynamics, expression of the nuclear M1F isoform significantly reduced replication fork progression in the absence of additional therapeutic manipulation (Fig. 4D). Importantly, mutation of M56 within the M1F construct restored replication fork progression, demonstrating that proper intracellular trafficking of CTSL is required for its ability to impair DNA replication.

### CTSL-dependent replication stress is coupled with impaired DNA repair capacity

To determine whether the therapy-induced replication defects observed by DNA fiber analysis translated into functional impairment of DNA repair, we performed GFP-based DNA repair reporter assays following treatment with CLF combined with ATR inhibitor. CLF/ AZD6738 combination reduced GFP reporter activity, indicating suppression of DNA repair competence under conditions that activate CTSL-dependent replication stress (Fig. 4E, F). Together, these findings indicate that CTSL-associated replication stress is not limited to altered fork dynamics, but is functionally coupled to impaired repair capacity, thereby lowering the threshold for irreversible fork collapse and cell death (Fig. 4). These findings establish that nuclear CTSL is not simply associated with replication stress but functionally restrains replication fork progression through coordinated remodeling of proteins governing replication licensing, replisome integrity, and DNA repair. Together with the preceding biochemical analyses, the DNA fiber studies demonstrate that CTSL-dependent proteolytic remodeling converts replication-competent ovarian cancer cells into a state characterized by persistent impairment of replication fork progression and loss of efficient DNA replication.

### CLF-based combinations create a therapeutically exploitable vulnerability in patient-derived ovarian cancer models

To determine whether CTSL-dependent remodeling of the replication machinery translates into clinically relevant therapeutic activity, we evaluated the efficacy of CLF-based combinations in primary ovarian cancer specimens established directly from malignant ascites. We reported patient derived primary ovarian ascitic (OVA) tumor cells were enriched based on PAX8 expression with minimal fibroblast contamination and multicellular spheroids were generated using autologous ascitic fluid to preserve the native tumor microenvironment[15].

Across fifteen independent patient-derived ascites cultures, CLF combined with the ATR inhibitor AZD6738 demonstrated broad and reproducible antitumor activity. Seven cultures exhibited very strong synergy with combination indices (CI) ranging from <0.1 to 0.5, while an additional seven cultures demonstrated moderate synergy with CI values between <0.7 and 0.8. Overall, 14 of 15 patient-derived cultures (93%) responded synergistically to CLF plus AZD6738, indicating that CTSL-directed therapeutic vulnerability is maintained across genetically heterogeneous primary ovarian cancers (Fig. 5A and B). Representative spheroid images from responsive samples demonstrated marked reductions in both spheroid number and spheroid size following combination treatment compared with either monotherapy (Figure S3). Quantitative analysis confirmed substantially greater inhibition of spheroid growth in highly responsive samples such as OVA-11 than in moderately responsive cultures including OVA-12, consistent with their respective combination indices. These findings demonstrate that CTSL-dependent therapeutic vulnerability extends beyond established ovarian cancer cell lines and is preserved in clinically relevant patient-derived three-dimensional cultures.

**Figure 5:**
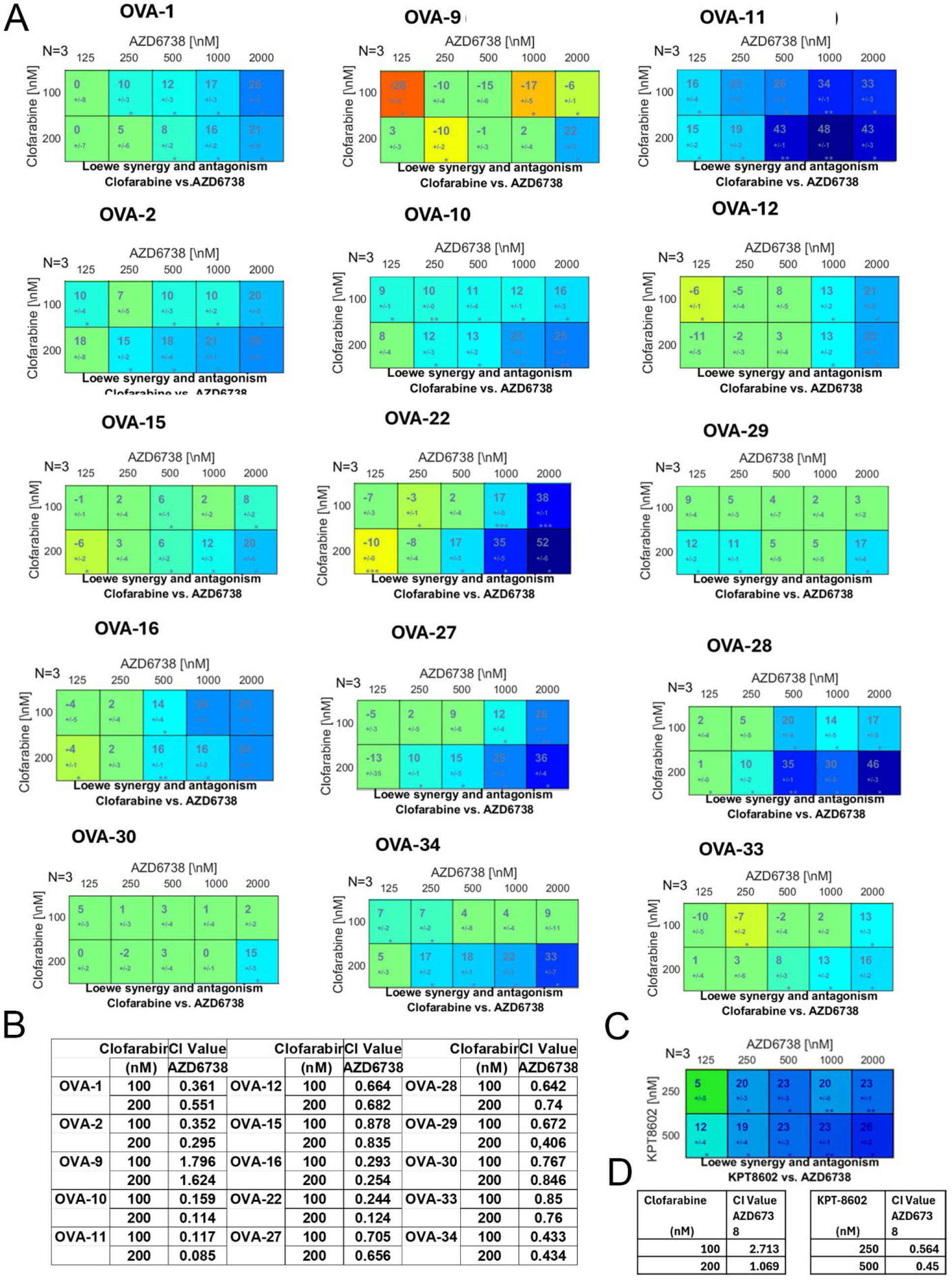
Evaluation of synergy upon treatment of CLF with AZD6738 in the ascites-derived ovarian cancer (OVA) cells *ex vivo*. **A)** Dual drug response assays for CLF + AZD6738 were performed in OVAs (OVA1-34) (n=15). Each assay included three technical replicates, repeated independently three times (N=3), and was analyzed using the Loewe synergy and antagonism matrix model with Combenefit software. The larger numeral in each box of the synergy matrix represents the synergy score, with negative values indicating antagonism. Green boxes indicate non-significant synergy scores, while colored boxes correspond to statistically significant results based on the synergism/antagonism scale, determined using a one-sample t-test. **B)** Combination index (CI) values across treatment panels were analyzed and presented in a table format. An average CI of 1 indicates an additive effect, CI < 1 represents synergy, and CI > 1 indicates antagonism. C and D) Similar dual response assays were performed for KPT8602 + AZD6738 combination in OVA9 *ex vivo* and combination index (CI) values calculated and presented in a table format.

### Restoration of nuclear CTSL trafficking overcomes acquired resistance to CLF-based therapy

Although the vast majority of patient-derived samples responded to CLF plus AZD6738, one culture (OVA-9) consistently exhibited resistance. Based on our previous studies, we hypothesized that this resistance reflected defective nuclear retention of CTSL resulting from failure of CLF to suppress the nuclear export receptor CRM1. Subcellular fractionation confirmed that CLF plus AZD6738 failed to promote sustained nuclear accumulation of CTSL in OVA-9 cells, consistent with loss of CTSL trafficking competence (data not shown). In contrast, direct pharmacologic inhibition of CRM1 using KPT8602 effectively restored nuclear retention of CTSL in these resistant cells. Replacement of CLF with KPT8602 therefore converted a resistant phenotype into a highly responsive state, with the KPT8602 plus AZD6738 combination producing marked synergy (CI <0.2) (Fig. 5C and D). BRCA and TP53 mutational status for all patient-derived samples was summarized[15]. These findings indicate that resistance to CLF-based therapy does not necessarily reflect loss of CTSL function but may instead result from impaired nuclear trafficking of CTSL. Restoration of nuclear trafficking competence therefore represents a potential strategy for overcoming acquired therapeutic resistance.

### CTSL-dependent remodeling of the replication machinery produces durable therapeutic responses in vivo

To determine whether CTSL-dependent remodeling of the replication machinery translates into therapeutic benefit *in vivo*, we evaluated CLF-AZD6738 combinations in the syngeneic KPCA with CCNE1 overexpression [21] and CCNE1 amplified PDX-PH115 ovarian cancer model. Mice bearing established intraperitoneal KPCA tumors were treated with CLF, AZD6738 as indicated (Fig. 6A). Whereas CLF, and AZD6738 monotherapies showed only mild to no significant effects on tumor growth, combining CLF with AZD6738 resulted in marked suppression of intraperitoneal tumor burden (Fig. 6B). These reductions translated into significant prolongation of overall survival, demonstrating durable therapeutic benefit following CTSL-directed combination therapy (Fig. 6C). To determine whether *in vivo* therapeutic responses were associated with activation of the CTSL-dependent replication program identified *in vitro*, endpoint tumor nuclear fractions were analyzed by immunoblotting. Tumors responding to combination therapy exhibited increased nuclear CTSL expression together with decreased CCNE1 and elevated γH2AX and phosphorylation of RPA32 at Ser33, indicating activation of replication-associated DNA damage signaling *in vivo* (Fig. 6D). On the other hand, the therapeutic efficacy of combining CLF with AZD6738 was evaluated using CCNE1-amplified PDX-PH115, a model previously identified as CLF-non-responsive *ex vivo* [15]. Contrary to the *ex vivo* findings, CLF monotherapy and the combination treatment reducing tumor burden towards baseline levels.in PH115-bearing mice compared to untreated controls (Fig. 6E, F). Moreover, Tumor area remained near baseline during the initial 4–5 weeks with the monotherapies, after which tumor growth increased above baseline. Following the 4-week treatment period, tumors continued to grow during the observation period across treatment groups. Furtherr, AZD6738 provided no additive benefit to CLF, as CLF monotherapy alone was able to reduce tumor progression to baseline (Fig. 6E, F). Importantly, these *in vivo* results diverged from the *ex vivo* PH115 model, where CLF monotherapy failed to induce nuclear CTSL localization, and only the combination treatment triggered nuclear CTSL alongside CCNE1 down-regulation (Figure S4). Nuclear fractionation of PH115 tumor tissues revealed that CLF monotherapy was indeed sufficient to drive nuclear translocation of CTSL *in vivo*. Furthermore, both CLF alone and the CLF/AZD6738 combination increased nuclear CTSL expression, down-regulated CCNE1, MCM3, and MCM6, and elevated γH2AX and p-RPA32 (Ser33) levels indicating activation of replication-associated DNA damage signaling *in vivo* (Fig. 6G). These molecular changes align with those observed in cultured ovarian cancer cells and other patient-derived models, confirming a shared mechanism of action across experimental systems.

**Figure 6:**
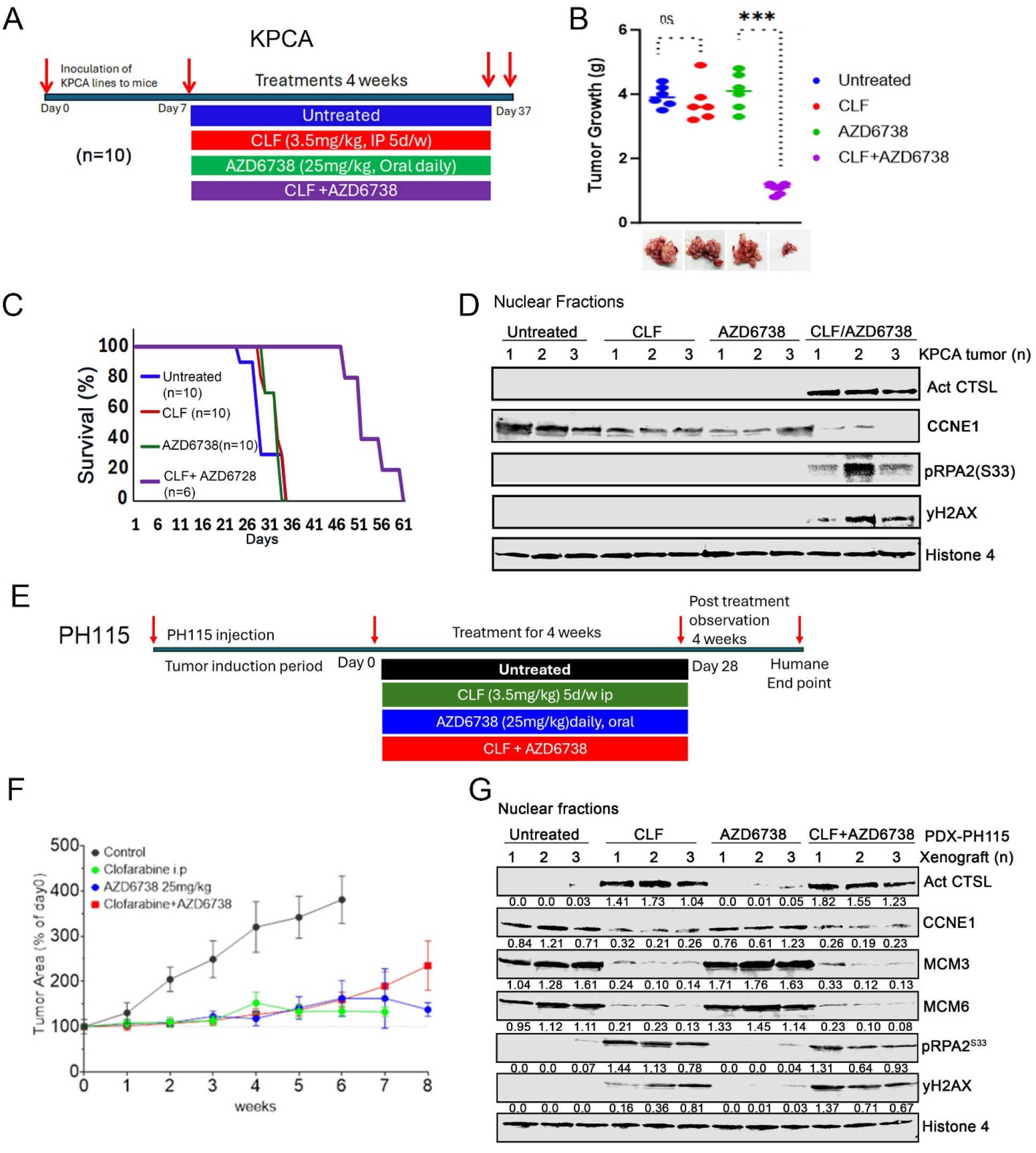
Antitumor efficacy of CLF and AZD6738 in the KPCA syngeneic tumor model and CCNE1 high PDX-PH115 model. Mice bearing KPCA tumors were treated with vehicle control, CLF, AZD6738, or the CLF/AZD6738 combination for 4 weeks, and tumor growth and survival were monitored. **A)** Schematic representation of treatment schedule for KPCA model **B)** Tumor growth was inhibited by CLF and AZD6738, with greater antitumor efficacy observed with the combination. **C)** Kaplan–Meier survival analysis with pairwise log-rank tests showed that CLF/AZD6738significantly improved survival compared with vehicle control or either monotherapy (**p < 0.01). **D)** Western blot analysis of nuclear fractions from KPCA tumor tissues was performed to assess nuclear CTSL, CCNE1, pRPA2 and γH2AX and the nuclear control Histone H3 levels. SCID mice bearing CCNE amplified PH115 tumors were treated with vehicle control, CLF, AZD6738, or the CLF/AZD6738 combination for 4 weeks, and tumor growth was monitored by ultrasound. **E)** Schematic representation of treatment schedule for PDX-PH115 model. **F)** Tumor growth over time, expressed as tumor area relative to baseline (day 0), as measured by ultrasound. Treatment was administered for 4 weeks, followed by an additional 4 week observation period **G)** Immunoblot analysis of nuclear-fractionated lysates from PDX-PH115 showing levels of nuclear CTSL, CCNE1, MCM3, MCM6, pRPA2 and γH2AX and the nuclear control Histone H3 levels. Statistics were analyzed by students t-test and significances were expressed as *\*p < 0.05, **p < 0.01 and ***p < 0.001*.

Collectively, these findings demonstrate that CTSL-dependent remodeling of the replication machinery generates a therapeutically exploitable vulnerability that extends from mechanistic studies in cultured ovarian cancer cells to patient-derived ascites spheroids and immunocompetent in vivo models. The concordance between molecular remodeling of the replication machinery, impaired replication fork progression, and durable antitumor activity supports a unified model in which therapeutic induction of nuclear CTSL converts replication-competent ovarian cancer cells into a state of replication collapse that can be effectively exploited by replication stress-directed combination therapy.

## DISCUSSION

Therapeutic resistance remains the principal obstacle to improving outcomes for patients with ovarian cancer despite the development of therapies that exploit replication stress and defects in homologous recombination. Although inhibition of ATR, CHK1, and PARP has demonstrated considerable promise, durable responses remain limited because tumor cells retain sufficient replication capacity and DNA repair function to survive therapeutic stress [5] [22]. The present study identifies nuclear cathepsin L (nCTSL) as a previously unrecognized regulator of this process. Rather than functioning solely as a lysosomal protease or mediator of DNA repair, our findings demonstrate that therapeutically induced nCTSL coordinately remodels the replication machinery by targeting proteins required for replication licensing, replisome integrity, and homologous recombination. This remodeling culminates in profound impairment of DNA replication, providing a mechanistic basis for the remarkable therapeutic synergy observed with CLF-based combinations.

Previous studies have largely viewed replication stress through the framework of checkpoint activation and DNA damage signaling [23, 24]. In this model, ATR, CHK1, and PARP function primarily to preserve replication fork stability and coordinate repair of damaged DNA [24–26]. Our findings extend this paradigm by demonstrating that therapeutic response is accompanied by coordinated proteolytic remodeling of the molecular machinery that sustains DNA replication itself. Rather than altering a single signaling pathway, nCTSL simultaneously reduces proteins governing replication licensing (CCNE1 and geminin), replisome function (MCM3 and MCM6), and homologous recombination (RAD51 and 53BP1) [15]. The simultaneous disruption of these interconnected processes provides a mechanistic explanation for why CLF-based combinations produce substantially greater biological effects than inhibition of individual replication stress pathways alone.

The functional consequences of this coordinated remodeling were demonstrated directly by DNA fiber analysis[27, 28]. Although checkpoint inhibition or nucleoside depletion individually produced only modest effects on DNA replication, combination treatment markedly impaired replication fork progression, an effect that persisted despite prolonged nucleotide labeling and was only partially rescued by exogenous deoxynucleosides (Fig. 4). These observations indicate that impaired DNA synthesis cannot be explained solely by depletion of nucleotide pools but instead reflects disruption of the molecular framework required to sustain efficient DNA replication. Consistent with this interpretation, CTSL depletion prevented these replication defects, whereas re-expression of the nuclear M1F CTSL isoform restored replication restraint, establishing nuclear CTSL as both necessary and sufficient for this phenotype. Additionally, the increased fork velocity observed following CTSL loss is consistent with our previous observations of reduced replication stress signaling and therapeutic resistance in CTSL-deficient cells, supporting a model in which CTSL functions as a regulator of replication homeostasis rather than merely a mediator of cell death.

Our findings further demonstrate that remodeling of the replication machinery is accompanied by progressive loss of homologous recombination capacity. Degradation of RAD51 and 53BP1 occurred[15] together with robust activation of γH2AX and pRPA2, indicating that cells entering this state experience persistent replication-associated DNA damage while simultaneously losing the capacity for efficient repair. Importantly, GFP-based DNA repair reporter assays revealed that CLF-based combinations suppress repair competence in a CTSL-dependent manner (Fig. 4). These findings indicate that CTSL-dependent replication stress is not restricted to altered fork dynamics but extends to functional inhibition of DNA repair pathways required for fork recovery and genome maintenance. Rather than representing independent molecular events, these observations support a model in which disruption of replication licensing, fork progression, checkpoint signaling, homologous recombination, and repair competence represents a coordinated cellular response initiated by therapeutic induction of nuclear CTSL. This integrated mechanism provides a biological explanation for the broad therapeutic activity observed following combinations of CLF with ATR, CHK1, or PARP inhibition (Fig. 1-6).

An important strength of the present study is the demonstration that this mechanism extends beyond established ovarian cancer cell lines. CLF-based combinations produced robust therapeutic responses in patient-derived ascites spheroids representing genetically heterogeneous recurrent ovarian cancers and demonstrated durable antitumor activity in immunocompetent syngeneic models. Moreover, resistance observed in one patient-derived culture was associated with defective nuclear retention of CTSL and could be overcome by pharmacologic inhibition of CRM1, restoring nuclear CTSL accumulation and therapeutic responsiveness (Fig. 5). These findings suggest that preservation of CTSL trafficking competence, rather than individual genomic alterations alone, may represent a previously unrecognized determinant of therapeutic response.

Although the present study establishes nuclear CTSL as a central regulator of replication machinery remodeling, several questions remain. The mechanisms governing substrate selection within the nucleus and the temporal sequence through which individual replication proteins are targeted remain to be fully defined. Likewise, although our findings demonstrate that restoration of nuclear CTSL is sufficient to remodel the replication machinery, additional studies will be required to determine how this process interfaces with chromatin organization, replication factory dynamics, and genome-wide replication programs. Addressing these questions will provide a more complete understanding of how proteolytic remodeling contributes to replication-associated therapeutic vulnerability.

Collectively, our findings support a model in which therapeutic induction of nuclear CTSL initiates a coordinated proteolytic remodeling program that dismantles multiple functional components of the replication machinery. Rather than acting solely through amplification of replication stress signaling, nCTSL shifts ovarian cancer cells from a state capable of sustaining productive DNA replication toward one characterized by impaired replication licensing, suppression of DNA repair competence, and persistent failure of replication fork progression. We propose that this transition represents a loss of replication competence, in which cells not only experience heightened replication stress but also lose the capacity to recover from replication-associated lesions. By simultaneously destabilizing replication forks and limiting repair capacity, nuclear CTSL lowers the threshold for irreversible fork collapse and promotes a state of replication catastrophe that is highly susceptible to therapeutic intervention. This framework unifies the molecular, functional, and translational findings presented here while providing a conceptual basis for exploiting CTSL-dependent vulnerabilities in ovarian cancer and potentially other replication stress-driven malignancies.

## MATERIALS AND METHODS

### Materials

All drugs, reagents, antibodies used are listed in Supplementary Table 1.

### Cell culture

All cell lines were cultured in their respective culture medium of DMEM/F12 or RPMI supplemented with 10% FBS and 1% penicillin/streptomycin in humidified incubators at 37°C and 5% CO_2_ atmosphere. Additionally, FT33 cells were generated through immortalization with human telomerase reverse transcriptase and SV40 large T antigen of non-malignant human fallopian tube secretory epithelial cells (FT33TAg). These cells were further transformed by either c-myc overexpression or by knockdown of p53 in combination with expression of mutant CDK4. Cells were treated with the indicated concentrations of control or clofarabine (CLF) with/without CHK1 inhibitor, AZD6738, CTSL inhibitor Z-FY-(tBu) DMK, or PARP inhibitor saruparib for 24 hours. For clonogenic assays, 300 cells per well were seeded in 24-well plates and treated for 10 days until colonies became visible; after which, they were stained using crystal violet dye and quantified using ImageJ software.

### Subcellular fractionation and analysis

Cells treated with the indicated concentrations of control or clofarabine (CLF) with/without CHK1 inhibitor, AZD6738 inhibitor, CTSL inhibitor, or PARP inhibitor saruparib in vitro and ex vivo were subjected to subcellular fractionation to isolate cytosolic, nuclear, and mitochondrial compartments using a NE-PER Kit and Mitochondrial Isolation Kit for Tissue (Thermo Fisher Scientific, USA) according to the manufacturer’s protocols. Briefly, the tissues/cells were washed, incubated in cytoplasmic extraction solution and then centrifuged. The supernatant served as the cytoplasmic fraction that was further processed using mitochondrial extraction buffer and centrifuged for the mitochondrial fractions in the pellet. The original residual pellet was processed using the nuclear extraction buffer and centrifuged to isolate the nuclear fractions. These fractions, along with whole-cell lysates, were analyzed by SDS–PAGE and served as adjuncts for further experiments.

### Generation of stable knockdown cell lines

PEO1 and PEO1-OLTRC4 (Gift from Dr. Xinyan Wu, Mayo Clinic), OVCAR4 cells cell lines were stably knocked down for CTSL using specific targeted shRNA from Sigma-Aldrich, following supplier’s protocol. CTSL-sh1 (GAATTGCCTCAGCTACTCTAA) CTSL-sh2 (TGCCTCAGCTACTCTAACATT) CTSL-sh3 (AGGCGATGCACAACAGATTAT) and with nontargeted control shRNA as control with Lipofectamine 3000 (Invitrogen, Carlsbad, CA) as per manufacturer protocol in the indicated cell lines. Similarly, PEO1 and PEO1-OLTRC4 cells were engineered for stable knockdown of ATR or ATM using shRNA constructs (ATR-AACCTCCGTGATGTTGCTTG; ATM-CCTTTCATTCAGCCTTTAGAA) purchased with corresponding non-targeting controls. Transfections were carried out and stable cell populations were selected with puromycin as previously described [29].

### Synergy assessment

Drug combination effects were evaluated using Combenefit software (http://sourceforge.net/projects/combenefit/), which compares the observed drug response surface to a theoretical reference model and generates a percentage synergy score for each point in the dose-response matrix. Synergy and antagonism distributions were visualized, and drug combinations of CLF with AZD6738 or prexasertib (n = 3) were specifically assessed using the Loewe additivity model. Statistically significant interactions were highlighted. One sample t-test was used to assess the significance of observed synergy scores [30].

### Homologous Recombination (HR) assay

OVCAR4 cells were stably transfected with the HR substrate pDR-GFP and transfected with pCβASceI plasmid for 24 hours and treated with indicated concentrations of CLF and AZD6738 alone and in combination for another 24 hours, and evaluated for the percent GFP positive fluorescence using flow cytometry.

### DNA fiber analysis

For the determination of tract length, ovcar4 cell pretreated CLF(5nM) and Prexasertib (5nM) for 10hs were labelled with 25μM CIdu for 20 min, after washing, cells were treated with 250 μM ICU and 1μM DN for 60 min. DNA Fiber spreads were prepared as described. After tyrosination and resuspension in ice-cold PBS, 2μl cells were diluted with 10 μl lysis buffer (200mM tris-HCL ph7.4, 50 mM EDTA, 0.5% SDS) on a glass slide then fixed in a methanol: acetic acid mixture (3:1) for 2 min at RT. Denaturing and blocking with 5% BSA+0.1% Tween 20, the slides were incubated in rat α-BrdU and mouse α-BrdU first antibodies for 2h. Fibers were treated with secondary antibodies for 1h at RT, allowed to air-dry and mounted in ProLong Gold antifade. Widefield pictures were taken with BioTek Lionneart microscopy. Data was evaluated and analysis by Graph Pad software.

### Immunoblot assays

Whole cell lysates and cellular fractions were processed using SDS-PAGE and analyzed by immunoblotting as reported earlier [29] using antibodies listed in Supplementary table 1. Secondary staining was achieved with secondary anti-mouse and rabbit with-680 or 800 IR dyes and scanned under the Odyssey Fc Imaging system (Bio-Rad, Hercules, CA). The normalized relative expression folds were calculated using ImageJ software.

### Immunofluorescence assay

The cells were treated with CLF with or without AZD6738/prexasertib at indicated conditions. Then, the cells were fixed and incubated using primary antibodies against p-RPA32, RPA32, yH2AX, CRM1 and CTSL (1:100) and incubated at 4°C overnight. Secondary staining was achieved with appropriate anti-mouse and rabbit Alexa fluor 488 or 595 secondary antibodies. Slides were mounted with Antifade reagent (Invitrogen, Carlsbad, CA), visualized using a Zeiss-LSM780 confocal microscope and corrected total cell fluorescence (CTCF) was obtained by ImageJ-Fiji software.

### Patient derived xenograft (PDX) models

Female SCID mice were intraperitoneally injected with primary ovarian tumor PH115 and allowed to establish PDX model. The mice were then randomized into 4 groups (n=10) and treated as; group 1: vehicle control group; group 2: CLF (3.5mg/kg, i.p. 5d/wk), group 3: AZD6738 (25mg/kg, oral daily); group 4: CLF+AZD6738 combination therapy. Mice were then treated for 4 weeks and observed for another 4 weeks, then sacrificed when tumor burden exceeded 10% of body weight in the control group. At the endpoint of this study, tumor burden, ascites accumulation if any, were assessed. All animal procedures were performed in accordance with protocols approved by the Mayo Clinic Institutional Animal Care and Use Committee (IACUC #A00006931).

### KPCA (Syngeneic) tumor model

The KPCA syngeneic tumor model was developed in C57BL/6 mice as described before [21]. In brief, three million cells were injected intraperitoneally and maintained for 7 days. The mice were then randomized into 4 groups (n=10), group 1: vehicle control; group 2: clofarabine (3.5 mg/kg 5d/w intraperitoneal injection) ; group 3: AZD6738 (Ceralasertib 25 mg/kg/day by oral gavage); group 4: combination of clofarabine and AZD6738; Mice were treated for 4 weeks and sacrificed when tumor burden surpassed 10% of initial body weight. Tumor parameters and survival of animals were measured. Pairwise log-rank tests were performed for survival analysis and statistical significance (*p < 0.05; **p < 0.01; ***p <0.001). Animal experiments were carried out under the approved protocols and guidelines of the Mayo Clinic Animal Care and Use Committee (IACUC number# A00007937).

### Statistical analysis

All experiments were performed in triplicates and independently repeated at least three times unless otherwise indicated. Data are presented as mean ± standard deviation (SD). Statistical significance (*p< 0.05; **p< 0.01; ***p< 0.001) was determined using Student’s t-test unless otherwise indicated. Statistical comparisons between groups for DNA fiber analysis were performed using Kruskal-Wallis test with Dunn’s post-hoc multiple comparisons test.

## Acknowledgements

We would also like to acknowledge the Mayo Ovarian SPORE for collection of OC ascites samples from patients and Dr. Ronny Drapkin (University of Pennsylvania Perelman School of Medicine) for the FT33(hTERT/TAg), FT33-shP53/CDK4^R24C^ and shPP2A-B56γ and FT33 c-Myc/TAg cells. We gratefully acknowledge contributions of the Mayo Clinic Cancer Center (P50 CA015803)-supported Pathology Research and Microscopy & Cell Analysis Cores.

## Contributions

Conceptualization: VS and PT. Methodology & Investigation: PT, LJ, SR, XH and JLV. Data assessment and verification: LJ, PT, JW, JC, JNB and VS. Supervision: VS. Writing—original draft: VS and PT. Writing—review & editing: PT, SR, KB, JC, SHK and VS. Correspondence and requests for materials should be addressed to VS.

## Data availability

Data is available within the Article or Supplementary Information.

## Ethics approval and consent to participate

All procedures involving animal were in accordance with the ethical standards of the ethics committee of Mayo Clinic Animal Care and Use Committee (IACUC number#A00006931). Clinical samples from patients were obtained at Mayo Clinic (IRB-approved protocol; 1288-03) and in association with the University of Minnesota Cancer Center Tissue Procurement Facility (IRB approval; 0702E01841).

## REFERENCES

1 da Costa AABA, Chowdhury D, Shapiro GI, D’Andrea AD, Konstantinopoulos PA. Targeting replication stress in cancer therapy. Nat Rev Drug Discov 2023; 22: 38–58.

2 Magdalou I, Lopez BS, Pasero P, Lambert SA. The causes of replication stress and their consequences on genome stability and cell fate. Semin Cell Dev Biol 2014; 30: 154–164.

3 Gorecki L, Andrs M, Korabecny J. Clinical Candidates Targeting the ATR-CHK1-WEE1 Axis in Cancer. Cancers (Basel) 2021; 13.

4 Guo YC, Wang J, Benedict B, Yang C, van Gemert F, Ma XH et al. Targeting CDC7 potentiates ATR-CHK1 signaling inhibition through induction of DNA replication stress in liver cancer. Genome Med 2021; 13.

5 Kim H, Xu H, George E, Hallberg D, Kumar S, Jagannathan V et al. Combining PARP with ATR inhibition overcomes PARP inhibitor and platinum resistance in ovarian cancer models. Nat Commun 2020; 11: 3726.

6 Berti M, Vindigni A. Replication stress: getting back on track. Nat Struct Mol Biol 2016; 23: 103–109.

7 Gaillard H, Garcia-Muse T, Aguilera A. Replication stress and cancer. Nat Rev Cancer 2015; 15: 276–289.

8 Ubhi T, Brown GW. Exploiting DNA Replication Stress for Cancer Treatment. Cancer Res 2019; 79: 1730–1739.

9 Goulet B, Baruch A, Moon NS, Poirier M, Sansregret LL, Erickson A et al. A cathepsin L isoform that is devoid of a signal peptide localizes to the nucleus in S phase and processes the CDP/Cux transcription factor. Mol Cell 2004; 14: 207–219.

10 Grotsky DA, Gonzalez-Suarez I, Novell A, Neumann MA, Yaddanapudi SC, Croke M et al. BRCA1 loss activates cathepsin L-mediated degradation of 53BP1 in breast cancer cells. J Cell Biol 2013; 200: 187–202.

11 Hiwasa T, Sakiyama S. Nuclear localization of procathepsin L/MEP in ras-transformed mouse fibroblasts. Cancer Lett 1996; 99: 87–91.

12 Duncan EM, Muratore-Schroeder TL, Cook RG, Garcia BA, Shabanowitz J, Hunt DF et al. Cathepsin L proteolytically processes histone H3 during mouse embryonic stem cell differentiation. Cell 2008; 135: 284–294.

13 Li Z, Peluffo G, Stevens LE, Qiu X, Seehawer M, Tawawalla A et al. KDM4C inhibition blocks tumor growth in basal breast cancer by promoting cathepsin L-mediated histone H3 cleavage. Nat Genet 2025; 57: 1463–1477.

14 Gonzalez-Suarez I, Redwood AB, Grotsky DA, Neumann MA, Cheng EH, Stewart CL et al. A new pathway that regulates 53BP1 stability implicates cathepsin L and vitamin D in DNA repair. Embo j 2011; 30: 3383–3396.

15 Thirusangu P, Jin L, Rao S, Ray U, Zhao A, Staub J et al. Drug-induced nuclear cathepsin L (nCTSL) defines a targetable DNA damage response axis and PARP inhibitor sensitivity in ovarian cancer. Cell Communication and Signaling 2026.

16 Karst AM, Jones PM, Vena N, Ligon AH, Liu JF, Hirsch MS et al. Cyclin E1 deregulation occurs early in secretory cell transformation to promote formation of fallopian tube-derived high-grade serous ovarian cancers. Cancer Res 2014; 74: 1141–1152.

17 Karst AM, Levanon K, Drapkin R. Modeling high-grade serous ovarian carcinogenesis from the fallopian tube. Proc Natl Acad Sci U S A 2011; 108: 7547–7552.

18 Ayhan A, Kuhn E, Wu RC, Ogawa H, Bahadirli-Talbott A, Mao TL et al. CCNE1 copy-number gain and overexpression identify ovarian clear cell carcinoma with a poor prognosis. Mod Pathol 2017; 30: 297–303.

19 Xu H, George E, Kinose Y, Kim H, Shah JB, Peake JD et al. CCNE1 copy number is a biomarker for response to combination WEE1-ATR inhibition in ovarian and endometrial cancer models. Cell Rep Med 2021; 2: 100394.

20 Xu H, George E, Gallo D, Medvedev S, Wang X, Kryczka R et al. Targeting CCNE1 amplified ovarian and endometrial cancers by combined inhibition of PKMYT1 and ATR. Res Sq 2024.

21 Iyer S, Zhang S, Yucel S, Horn H, Smith SG, Reinhardt F et al. Genetically Defined Syngeneic Mouse Models of Ovarian Cancer as Tools for the Discovery of Combination Immunotherapy. Cancer Discov 2021; 11: 384–407.

22 O’Connor MJ. Targeting the DNA Damage Response in Cancer. Mol Cell 2015; 60: 547–560.

23 Forment JV, O’Connor MJ. Targeting the replication stress response in cancer. Pharmacol Ther 2018; 188: 155–167.

24 Smith HL, Willmore E, Prendergast L, Curtin NJ. ATR, CHK1 and WEE1 inhibitors cause homologous recombination repair deficiency to induce synthetic lethality with PARP inhibitors. Br J Cancer 2024; 131: 905–917.

25 Dobbelstein M, Sørensen CS. Exploiting replicative stress to treat cancer. Nat Rev Drug Discov 2015; 14: 405–423.

26 Zeman MK, Cimprich KA. Causes and consequences of replication stress. Nat Cell Biol 2014; 16: 2–9.

27 Ray Chaudhuri A, Callen E, Ding X, Gogola E, Duarte AA, Lee JE et al. Replication fork stability confers chemoresistance in BRCA-deficient cells. Nature 2016; 535: 382–387.

28 Quinet A, Tirman S, Jackson J, Šviković S, Lemaçon D, Carvajal-Maldonado D et al. PRIMPOL-Mediated Adaptive Response Suppresses Replication Fork Reversal in BRCA-Deficient Cells. Mol Cell 2020; 77: 461–474.e469.

29 Jung D, Khurana A, Roy D, Kalogera E, Bakkum-Gamez J, Chien J et al. Quinacrine upregulates p21/p27 independent of p53 through autophagy-mediated downregulation of p62-Skp2 axis in ovarian cancer. Sci Rep 2018; 8: 2487.

30 Chou TC. Drug combination studies and their synergy quantification using the Chou-Talalay method. Cancer Res 2010; 70: 440–446.

