## Supplemental material for "Nuclear Cathepsin L Remodels the Replication Machinery to Create a Therapeutic Vulnerability in Ovarian Cancer"

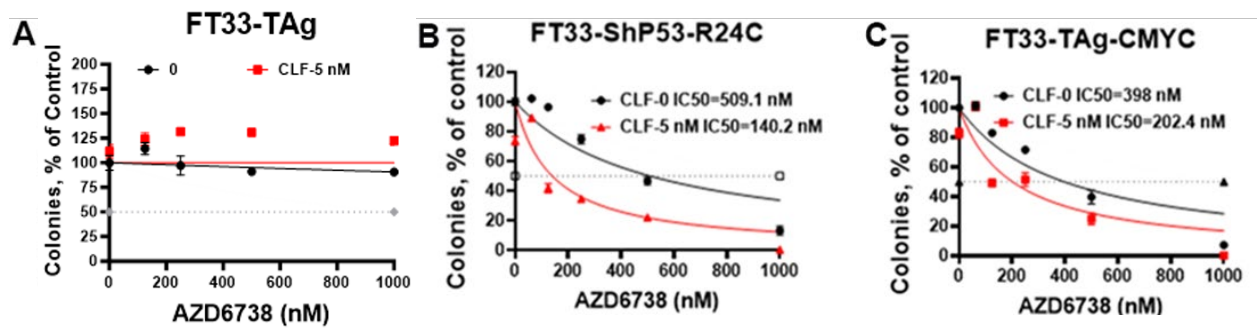

**Figure S1. CLF selectively enhances sensitivity to ATR inhibition in transformed fallopian tube secretory epithelial cells.** Colony-formation assays were performed in (A) immortalized FT33-TAg cells, (B) FT33 cells expressing shRNA targeting *TP53* and the *TP53* R24C mutant (FT33-ShP53-R24C), and (C) FT33 cells expressing SV40 large T antigen and *CMYC* (FT33-TAg-CMYC). Cells were treated with increasing concentrations of the ATR inhibitor (AZD6738; 0–1,000 nM) alone (CLF 0; black) or in combination with 5 nM CLF (red). Colony formation is expressed as the percentage of vehicle-treated control. Dashed horizontal lines indicate 50% colony formation. CLF had minimal effect on the response of immortalized FT33-TAg cells to AZD6738, whereas it enhanced AZD6738 mediated suppression of colony formation in transformed FT33-ShP53-R24C and FT33-TAg-CMYC cells.

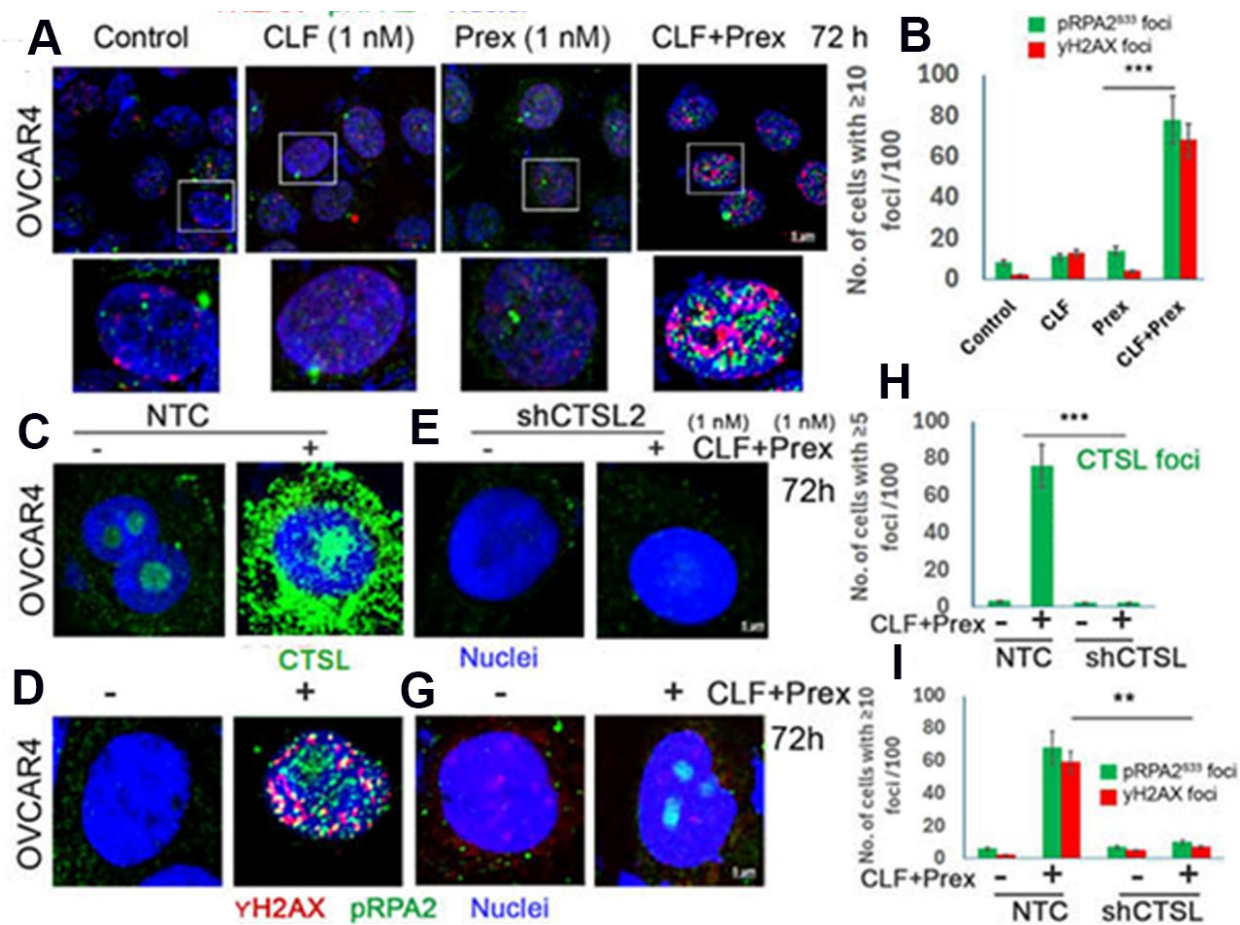

**Figure S2. CTSL depletion suppresses CLF/prexasertib-induced replication-stress signaling in OVCAR4 cells.** (A–B) OVCAR4 cells were treated with CLF (1 nM), prexasertib (Prex; 1 nM), or both for 72 h and stained for pRPA2<sup>S33</sup> (green), γH2AX (red), and nuclei (DAPI; blue). Combined treatment increased pRPA2<sup>S33</sup> and γH2AX foci. (C–I) Nontargeting control (NTC) or CTSL-depleted (shCTSL) OVCAR4 cells were treated with CLF plus prexasertib for 72 h. CTSL depletion reduced nuclear CTSL foci and prevented induction of pRPA2<sup>S33</sup> and γH2AX foci. Data are presented as the mean ± SD from three independent experiments. $P < 0.01$; \*$P < 0.001$.

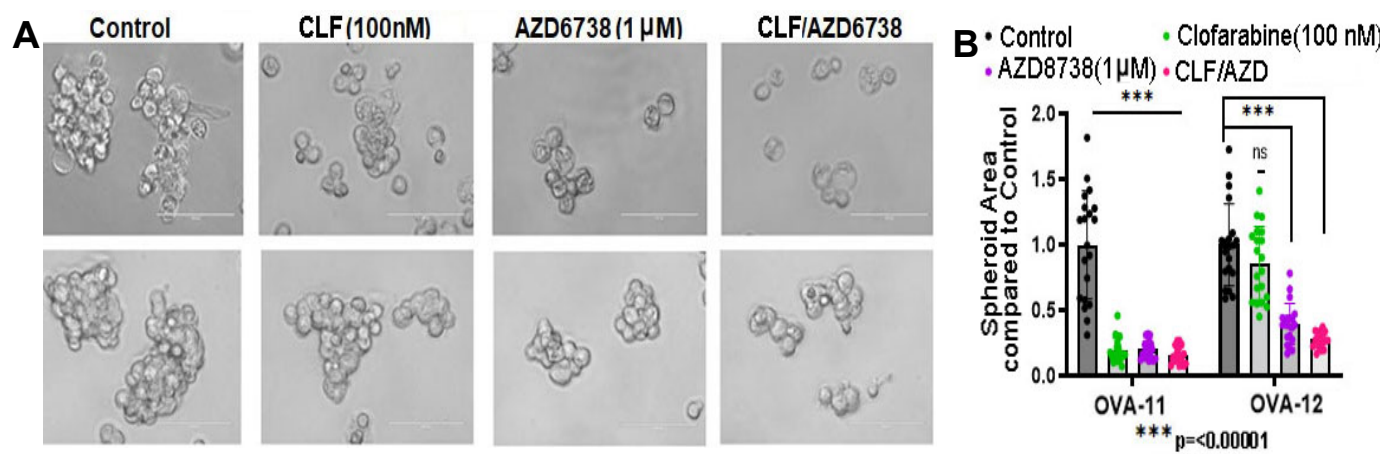

**Figure S3. Combined CLF and AZD6738 treatment suppress patient-derived ovarian cancer spheroid growth.** (A) Representative images of OVA-11 and OVA-12 spheroids treated with vehicle control, clofarabine (CLF; 100 nM), AZD6738 (1 μM), or the CLF/AZD6738 combination. Combination treatment reduced both spheroid number and size compared with either alone. (B) Combined CLF/AZD6738 treatment produced greater suppression of spheroid growth in the highly responsive OVA-11 spheroid cells than in the moderately responsive OVA-12. \*\*\*P < 0.001; ns, not significant.

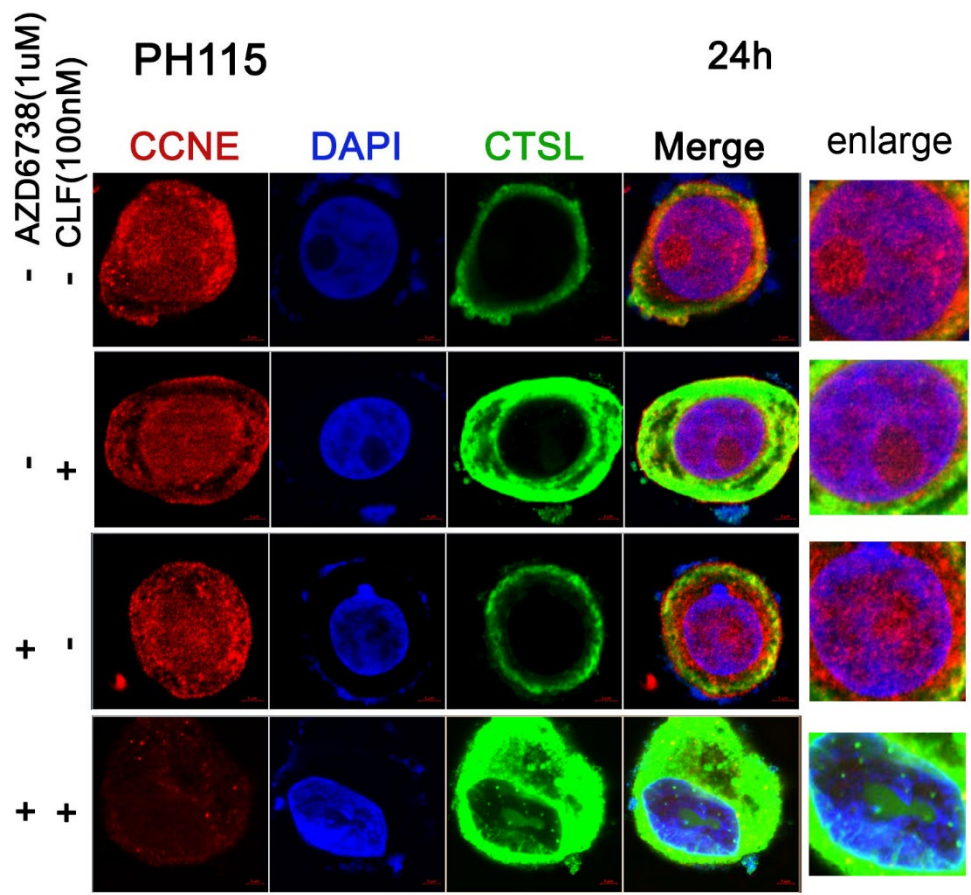

**Figure S4. CLF and AZD6738 combination treatment induce nuclear CTSL in PH115.** *Ex vivo* culture of PH115 treated with CLF and AZD6738 exhibited nuclear CTSL with decreased CCNE1.

**Supplementary Table 1**

| **Sl no** | **Name of the chemicals/reagent/plasmid** | **Catalog number** | **RRIDs** | **Manufacturer** |
| --- | --- | --- | --- | --- |
| 1 | Clofarabine | HY-A0005/CS-0373 |  | MedChemExpress (Monmouth Jct., NJ) |
| 2 | AZD6738 / ceralasertib | HY-19323 |  | MedChemExpress (Monmouth Jct., NJ) |
| 3 | Prexasertib | HY-18174 |  | MedChemExpress (Monmouth Jct., NJ) |
| 4 | Saruparib | HY-132167 |  | MedChemExpress (Monmouth Jct., NJ) |
| 5 | Deoxynucleosides cocktail | D8668 |  | Sigma |
| 6 | CldU | C6891 |  | Sigma |
| 7 | IdU | 2100357.2 |  | MP Biomedicals |
| 8 | Z-FY(tBU)-DMK | sc-222423 |  | Santa Cruz Biotechnology (Dallas, TX) |
| 9 | Eltanexor (KPT-8602) | S8397 |  | Selleckchem (Houston, TX) |
| 10 | cell lysis buffer | #9803S |  | Cell Signaling Technology |
| **Sl no** | **Name of the Antibodies/Plasmid/Kits** | **Catalog number** | **RRIDs** | **Manufacturer** |
| 1 | Anti-CRM1 | #46249S | AB_2799298 | Cell Signaling Technology |
| 2 | anti- PAX8 | sc-81353 | AB_1127048 | Santa Cruz |
| 3 | anti-GAPDH | sc-47724 | AB_627678 | Santa Cruz |
| 4 | anti-C9 tag | sc-390000 | AB_2894830 | Santa Cruz |
| 5 | anti-KPNB1 | sc-137016 | AB_2133993 | Santa Cruz |
| 6 | anti-β actin | sc-47778 | AB_626632 | Santa Cruz |
| 7 | anti- $\alpha$ tubulin | sc-5286 | AB_628411 | Santa Cruz |
| 8 | anti-53BP1 | #11940 | AB_2637071 | Abcam |
| 9 | anti- RAD51 | #ab133534 | AB_2722613 | Abcam |
| 10 | Anti-CTSL | #10486-R221 | SCR_003697 | Sino Biological |
| 11 | anti- Flag tag | #9291S | AB_10950495 | Abcam |
| 12 | anti-hFAP | # AF3715 | AB_2102369 | R & D systems |
| 13 | Anti-Cyclin E1 | sc-245 |  | Santa Cruz |
| 14 | Anti-pRPA2/RPA32 Ser33 | sc-48425 |  | Santa Cruz |
| 15 | Anti-γH2AX | GTX108272 |  | Genetex |
| 16 | Anti-Histone H3 | 9727S |  | Cell Signaling Technology |
| 17 | Anti-MCM3 | 15597-1-AP |  | Proteintech |
| 18 | Anti-MCM6 | 13347-2-AP |  | Proteintech |
| 19 | Anti-Geminin / GMNN | 52508 |  | Cell Signaling Technology |
| 20 | Anti-pCHK1 Ser345 | 2348S |  | Cell Signaling Technology |
| 21 | Anti-CHK1 | sc-8408 |  | Santa Cruz |
| 22 | shCTSL -TRC1.5 plasmid | SHCLNG-NM 001912 |  | Sigma Aldrich |
| 23 | pcDNA3.1-hCathepsin L construct | #11250 | Addgene_11250 | Addgene |
| 24 | NE-PER nuclear and CE-PER | #78833 |  | Life Technologies |
| 25 | IRDye 680 LT goat anti mouse IgG, red | 926-68020 | AB_2687826 | Li-Cor, Lincoln, NE, U.S.A. |
| 26 | IRDye 800CW Donkey anti-Mouse IgG, green | 926-32212 | AB_2716622 | Li-Cor, Lincoln, NE, U.S.A. |
| 27 | IRDye 800 goat anti rabbit IgG, green, | 926-32211 | AB_621843 | Li-Cor, Lincoln, NE, U.S.A. |
| 28 | IRDye 680RD Donkey anti-Rabbit IgG, red | 926-68073 | AB_10954442 | Li-Cor, Lincoln, NE, U.S.A. |
| 29 | Mitochondrial Isolation Kit | ab65320 |  | Abcam |
| 30 | anti-BrdU for CldU | ab6326 |  | Abcam |
| 31 | anti-BrdU for IdU | 347580 |  | BD Biosciences |
| 32 | Puromycin | A11138-03 |  | Gibco |
| 33 | RPA32 / total RPA32 antibody | 35869 |  | Cell Signaling Technology |
